# Assessment of the impact of shared data on the scientific literature

**DOI:** 10.1101/183814

**Authors:** Michael P. Milham, R. Cameron Craddock, Michael Fleischmann, Jake Son, Jon Clucas, Helen Xu, Bonhwang Koo, Anirudh Krishnakumar, Bharat B. Biswal, F. Xavier Castellanos, Stan Colcombe, Adriana Di Martino, Xi-Nian Zuo, Arno Klein

## Abstract

Data sharing is increasingly recommended as a means of accelerating science by facilitating collaboration, transparency, and reproducibility. While few oppose data sharing philosophically, a range of barriers deter most researchers from implementing it in practice (e.g., workforce and infrastructural demands, sociocultural and privacy concerns, lack of standardization). To justify the significant effort required for sharing data (e.g., organization, curation, distribution), funding agencies, institutions, and investigators need clear evidence of benefit. Here, using the International Neuroimaging Data-sharing Initiative, we present a brain imaging case study that provides direct evidence of the impact of open sharing on data use and resulting publications over a seven-year period (2010-2017). We dispel the myth that scientific findings using shared data cannot be published in high-impact journals and demonstrate rapid growth in the publication of such journal articles, scholarly theses, and conference proceedings. In contrast to commonly used ‘pay to play’ models, we demonstrate that openly shared data can increase the scale (i.e., sample size) of scientific studies conducted by data contributors, and can recruit scientists from a broader range of disciplines. These findings suggest the transformative power of data sharing for accelerating science and underscore the need for the scientific ecosystem to embrace the challenge of implementing data sharing universally.

---

Now more than ever, the potential and actual benefits of open data sharing are being debated in the pages of premier scientific journals, funding agency communications, scientific meetings and workshops^1,2,3^. Throughout these discussions an array of potential benefits are acknowledged, ranging from increased transparency of research and reproducibility of findings to decreased redundancy of effort and the generation of large-scale data repositories that can be used to achieve more appropriate sample sizes for analyses. Equally important, data sharing is commonly described as a means of facilitating collaboration across the broader scientific community.

Despite its potential, for many, the benefits of data sharing are more theoretical than practical^2,4^. The reality is that data sharing is relatively limited in many disciplines and little information on its outcomes exists^5^. In the absence of clear demonstrations of data sharing’s impact, debates on the topic are dominated by its formidable — albeit hypothetical — downsides. Common concerns include loss of competitive advantage (especially for junior investigators)^6^, fear of being scooped with one's own data, scientifically unsound uses of the data, and concerns that high-impact journals will not accept manuscripts that report findings generated by secondary analysis of open datasets.

To assess the tangible benefits of open data sharing, we provide a bibliometric analysis of a large brain image data sharing initiative. The brain imaging community is a particularly valuable target for examination, as its challenges are representative of those commonly encountered in biomedical research. The high costs and workforce demands required to capture primary data limit the ability of individual labs to generate properly powered sample sizes. These obstacles are amplified when addressing more challenging (e.g., developing, aging, clinical) populations or attempting biomarker discovery — both prerequisites for achieving clinically useful applications. Inspired by the momentum of molecular genetics, the first functional neuroimaging data sharing initiative was launched in 2000^7^, though it encountered logistical challenges (e.g., lack of standardization for task-based fMRI methods) and vigorous social resistance. Since then, a range of initiatives for sharing brain imaging data have emerged (e.g., OASIS^8^, ADNI^9^, Human Connectome Project^10^, OpenfMRI^11^).

While some open data sharing initiatives work to aggregate and share previously collected datasets, others explicitly generate large-scale data resources for the purposes of sharing. The present work focuses on the International Neuroimaging Data-sharing Initiative (INDI)^12^, as it uniquely embodies both of these models of sharing. The bibliometric measures we have employed could be easily applied to other initiatives in future work. Another distinctive aspect of INDI is its reliance on the formation of grassroots consortia as a primary vehicle for achieving its goal of aggregating and sharing previously collected data. Self-initiated and organized by scientists in the community, these consortia aggregate and share independently collected data from sites around the world. Examples of INDI-based consortia include the 1000 Functional Connectomes Project (FCP; n = 1,414, released in December, 2009)^13^, the ADHD-200 (n = 874, released in March, 2010)^14^, the Autism Brain Imaging Data Exchange (ABIDE; n = 1,112, released in August, 2012)^15^ and the Consortium for Reliability and Reproducibility (CoRR; n = 1,629, released in June, 2014)^16^. In the present work, we use the grassroots consortium component of the INDI model to examine the relative benefits of open data sharing versus ‘pay to play’ models, in which only those who give data can benefit from sharing. To examine the benefits of data resources explicitly generated for the purposes of sharing, we use INDI’s Nathan Kline Institute-Rockland Sample (NKI-RS) initiatives, a combination of large-scale crosssectional and longitudinal multimodal imaging samples of brain development, maturation and aging (ages 6.0-85.0)^17,18^ (initial release in October, 2010; quarterly releases ongoing, current n = 1,000+). INDI efforts have been lauded by funding agencies, journal editors, and members of the imaging community. However, such subjective recognition does not quantify research impact. Drawing from the field of bibliometrics, we carried out a range of citation analyses^19^ to quantify the impact of INDI datasets on the brain imaging and broader scientific literature.

We started our bibliometric analysis with a search for publications that used INDI datasets. This was a non-trivial task due to the lack of requirements for author-line recognition of INDI, a policy intended to maximize freedom of use for the data. We identified publications using a full-text search in Google Scholar; the following names and URLs were included as keywords: ‘fcon_1000.projects.nitrc.org’, ‘Rockland Sample’, ‘1000 Functional Connectomes’, ‘International Neuroimaging Data-Sharing Initiative’, ‘Autism Brain Imaging Data Exchange’, ‘ADHD-200’, and ‘Consortium for Reproducibility and Reliability’. Next, we downloaded all available PDF files for manual review by a team of five research assistants, who classified each as *‘downloaded and used INDI subject data’, ‘only mentions or references INDI data’, ‘used INDI scripts but not INDI data’*, or *‘irrelevant’*. To facilitate this process and enable rapid review, each PDF was converted to a text file (using the Unix-based *pdftotext* shell command). Paragraphs including the keywords from the Google Scholar search were then identified and extracted from each PDF for review in an automated fashion using regular expressions (code for analyses is available at https://github.com/ChildMindInstitute/BiblioReader/blob/165ddc56779a5e55149184a0f95b7c14874cf0c5/biblioreader/texttools/texttools.py); full PDFs were available to the reviewers for verification. Classifications were determined from the consensus of two independent reviewers; conflicts were resolved by a third. During this step, research assistants also indicated the type of publication (e.g., thesis, book chapter, peer-reviewed journal article, non peer-reviewed journal article, preprint) for each paper.

## Data Use

Our keyword-based search identified 1,541 possible INDI-related publications as of March 22, 2017, of which 913 were determined to have used data from INDI. Figure 1a provides a non-cumulative breakdown of the 913 publications by year and initiative, revealing steady yearly increases in shared data use. Author affiliations for the 913 publications using INDI data spanned 50 countries across 6 continents, with peak affiliation densities in the United States (48.5%), China (10.7%), Germany (6.5%), and the United Kingdom (6.0%) (see http://fcon1000.projects.nitrc.org/indi/bibliometrics/map/map.html for the world map of author affiliations generated using the Google Maps JavaScript API V3). The overwhelming majority of publications were either peer-reviewed journal articles (n = 739; 81%) or preprints (n = 65; 7%); scholarly theses had a substantial presence as well (n = 58 [33 doctoral/19 master's/4 bachelor's/2 unspecified]), demonstrating the potential value of shared data for trainees and early career investigators (see Figure 1b). As would be expected given the brain imaging focus of the INDI consortia, the relative majority (45.7%) of publications were in journals focused on general neuroscience, neurology, psychiatry, and psychology. However, there was evidence of INDI datasets penetrating other domains as well (e.g., mathematics, computer science, physics, and engineering journals accounted for 6.6% of publications) (see Figure 1c).

**Figure 1.**
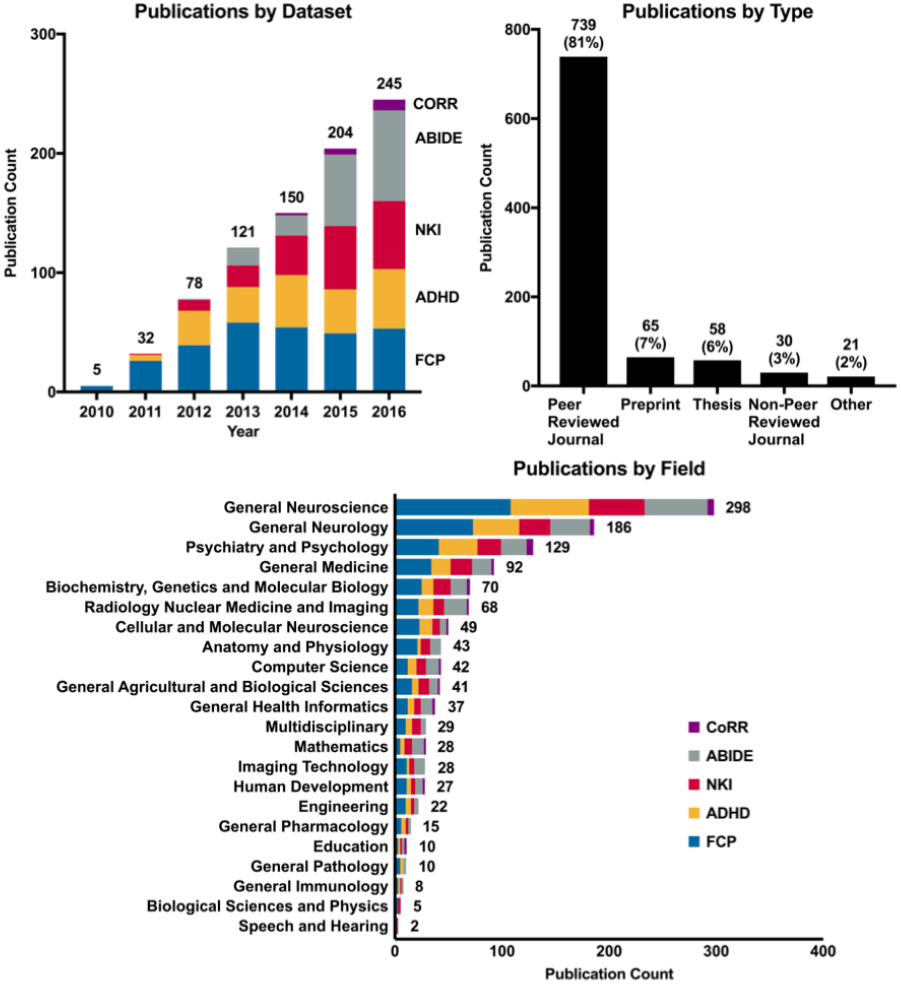
Publications that used INDI shared data. **a-c**, Publications sorted by INDI dataset and year, for the period of 2010 to 2016 (publications from 2017 not depicted as year was in progress at time of present work) (**a**), publication type (**b**), and journal discipline (limited to peer-reviewed publications; based on Web of Science classifications) (**c**).

## Publication Impact

The impact of each of the major INDI efforts (FCP, NKI-RS, ADHD-200, ABIDE, CoRR) on the scientific literature was quantified using an array of commonly used citation-based indices (see Table 1). As of March 22, 2017, the 913 publications that explicitly used INDI data had been cited 20,697 times by publications referenced in Google Scholar, with an average of 4.4 citations per article per year; h-indices for the five initiatives ranged from 7 to 52 (overall: 66) and i10-indices from 6 to 123 (overall: 295). The FCP and ADHD-200 have had the highest impact to date across various measures, though this likely reflects their older age compared to other initiatives (e.g., ABIDE, NKI-RS), which enjoy greater publication growth in recent years (see Figure 1a).

**Table 1.** Quantifying Impact of INDI Efforts Using Common Publication-based Indices.

| Initiative | Total Citations | Mean Citations Per Year | Mean Total Citations | h-Index | h-Index (5-year) | i10 index | i10-Index (5-year) |
| --- | --- | --- | --- | --- | --- | --- | --- |
| FCP | 13147 | 7.3 ± 20.8 | 40.8 ± 140.4 | 52 | 43 | 123 | 104 |
| ADHD-200 | 2935 | 2.9 ± 5.3 | 14.5 ± 36.0 | 33 | 31 | 67 | 66 |
| ABIDE | 1875 | 2.5 ± 6.9 | 9.2 ± 28.3 | 22 | 22 | 44 | 44 |
| CoRR | 357 | 4.1 ± 7.5 | 16.3 ± 34.6 | 7 | 7 | 6 | 6 |
| NKI-RS | 2383 | 3.3 ± 5.6 | 11.9 ± 24.7 | 29 | 29 | 55 | 54 |
| Total | 20697 | 4.4 ± 11.7 | 20.4 ± 72.9 | 66 | 58 | 295 | 274 |

To address concerns about publishing analyses of shared data in high impact journals, we also examined journal impact factors for publications using INDI data. While the assessment of journal impact remains somewhat controversial given the growing number of indices available (e.g., impact factor, CiteScore, altmetrics)^20^, several articles that used INDI data have been published in high-impact specialized journals (e.g., *Biological Psychiatry*, *Neuron*) and general interest journals (e.g., *Proceedings of the National Academy of Sciences, Nature Communications*) (see Figure 2a for the 15 highest impact journals in which publications using INDI datasets have appeared based on CiteScore^20^). As shown in Figure 2b, of all publications measured by 2015 CiteScore values, 50 percent were published in a journal with a CiteScore of 4.05 or higher, 25 percent with a CiteScore of 6.71 or higher, 5 percent with a score of 8.84 or higher and 1 percent with a score of 12.02 or higher. Two of the three journals with the highest number of publications were *NeuroImage* and *Human Brain Mapping*, which are among the highest ranked field-specific brain imaging journals.

**Figure 2.**
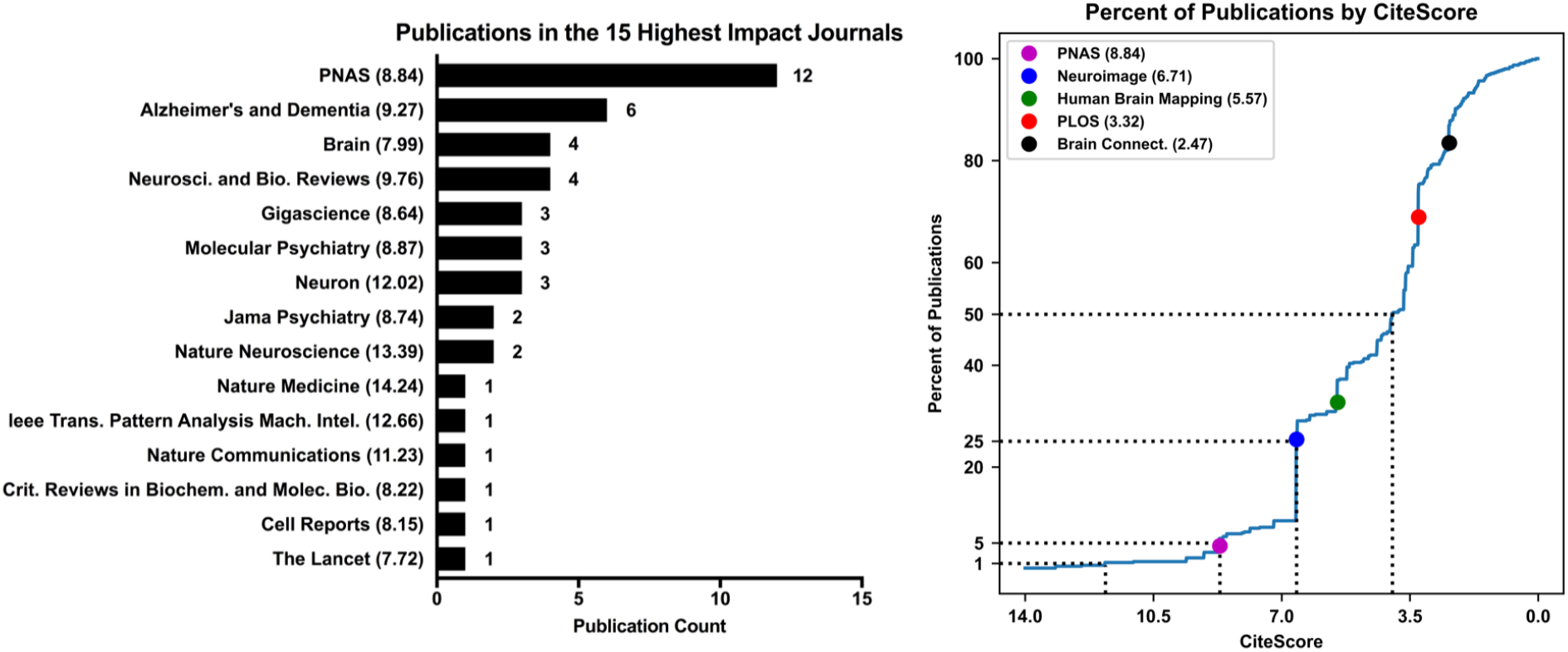
Quantification of Publication Impact. **a**, 15 highest impact journals in which publications using INDI datasets have appeared (based on CiteScore; number of publications depicted for each journal). **b**, Cumulative density function for CiteScores of publications based on INDI data. Select publications depicted to provide reference points to assist in interpretation of CiteScore.

## Beneficiaries of Sharing

A common alternative to open data sharing is the “pay to play” model, where only those who contribute data can gain access to shared data. While such models can incentivize data sharing, they miss out on valuable analyses that researchers lacking data to contribute would perform if given the opportunity. INDI’s consortium model provides a unique opportunity to compare use of shared data by contributing and non-contributing researchers. Specifically, for each initiative (FCP, ADHD-200, ABIDE, CoRR, NKI-RS), “contributing authors” were defined as any co-author of the announcement publication for the respective initiative. Using this definition, 90.3% of INDI-based publications were authored by research teams that did not include any data contributors. As shown in Figure 3, the number of publications by non-contributors is rapidly increasing year to year. This differential between publications authored by contributors vs. non-contributors reflects the potential missed opportunity associated with the ‘pay to play’ model of data sharing.

**Figure 3.**
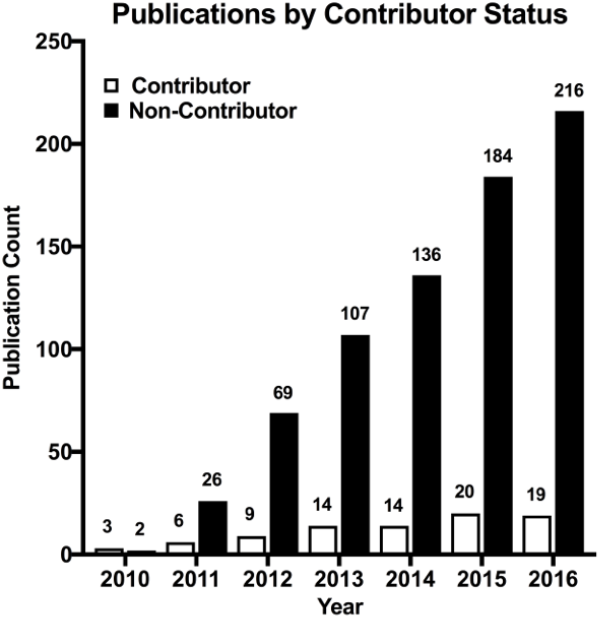
Data use by authors. Breakdown of publications by contributor status, for the period of 2010 to 2016 (publications from 2017 not depicted as year was in progress at time of present work).

The publication patterns of INDI consortia members can also be used to glean insights into the benefits of contributing data beyond inclusion as a coauthor on data announcement or descriptor papers. To accomplish this, we focused on the ADHD-200 and ABIDE consortia, as they consist of data from clinical populations, which are among the most costly to generate. For each ADHD-200 or ABIDE paper co-authored by a data contributor, we calculated the difference between the amount of data used in the publication (i.e., sample size) and the total contribution to the consortium (from co-authors on the manuscript). The median difference between publication sample size and data contribution by coauthors was 286 for the ADHD-200 and 142.5 for ABIDE. Obtaining a similar increase in sample size by acquiring data from these clinical populations in a single lab would have been expensive and time consuming. Interestingly, we found that 20% of INDI-based publications from ABIDE data contributors used fewer samples than they contributed — largely reflecting more restrictive inclusion criteria related to age, sex, diagnostic phenotype, and/or image quality. Application of such criteria was made possible by the availability of shared data, and presumably enhanced the validity and reproducibility of the findings.

Another means by which shared datasets are becoming increasingly available is the resource generation model, in which data are specifically collected for the purpose of sharing (e.g., Human Connectome Project, Brain Genomics Superstruct, NIH ABCD, Child Mind Institute Healthy Brain Network). This model is advantageous in that the explicit open access intent allows researchers and funding agencies to justify investing in the creation of data resources that are notably larger in scale and broader in scope than what would typically be acquired by a single team. In INDI, the NKI-RS is an ongoing coordinated effort of three principal investigators and four funded projects dedicated to generating an open lifespan data resource for the scientific community. To date, 189 articles have been published based on NKI-RS data, 167 of which did not include the NKI-RS PIs and 76 of which were written by individuals completely outside of their publication sphere (i.e., there is no detectable relationship between the PIs and the authors of these papers based on coauthorship histories in the literature).

## Impact Beyond

It is important to note the various forms of impact that are not captured by the present analyses. First, our searches revealed 639 publications that mentioned INDI in their text but did not use INDI data, suggesting that INDI and resultant research has impacted the thinking of authors in ways other than simply providing data. Second, 71 publications employed either the scripts used for the analysis of data in the initial FCP release manuscript^13^ or their derivative platforms (^21,22^). Third, INDI has also given rise to the Preprocessed Connectomes Project (http://preprocessed-connectomes-project.org), which shares processed forms of INDI data, as well as quality measures. Finally, it is worth acknowledging the unpublished training and testing of protocols and algorithms not captured by our analyses, as well as the unique scientific opportunities created by the aggregation of independently collected datasets. Examples of such opportunities include the ability to demonstrate reproducibility of findings across independent datasets (e.g., ^23^), to assess potential solutions for overcoming batch effects (i.e., scanner/protocol differences) (e.g., ^24-26^), and to provide robust assessments of statistical noise (e.g., ^27^).

## Concluding Remarks

Having demonstrated the impact of data sharing, the remaining challenge is to make sharing a widespread reality. Multiple advances are necessary to accomplish this change. First, there is a need for greater incentivization of both sharing and using open datasets. While funding agencies increasingly espouse mandates to share data and encourage secondary data analysis, voices vilifying openly shared data and its users^28^ continue to be expressed in top journals. Second, the mechanisms for recognizing data sharing contributions remain underspecified in, for example, grant, promotion, or tenure reviews^29^. Third, widespread data sharing requires infrastructure. To date, the storage costs of the INDI have been relatively limited, requiring about 10TB to share over 15,000 datasets. However, resource limitations must be considered as data sharing and the size of shared datasets continue to grow along with acceptance of data sharing and of data sharing mandates by funding agencies and journals. Central to these considerations will be decisions regarding the emphasis to be placed on centralized vs. federated models for data storage, as the scope and scale of data sharing increases. Such decisions are non-trivial, as they entail a range of financial, logistical and ethical questions regarding data maintenance and privacy. Finally, increased harmonization of data acquisition procedures is needed within each scientific field. Adoption of common strategies for phenotyping (e.g., common data elements) can dramatically improve the value of shared datasets. Similarly, the harmonization of data acquisition procedures (e.g., MRI scan sequences, experimental procedures, phenotyping instruments) and establishing quality assessment standards will further improve the quality and value of data sharing (see ^30^ for a comprehensive discussion related to brain imaging). We assert that it is the responsibility of the entire scientific ecosystem, from funding agencies to junior scientists, to accelerate the pace of progress by making data sharing the norm.

## Acknowledgements

The Child Mind Institute provides primary funding for the INDI team, with additional support provided by the Nathan S. Kline Institute for Psychiatric Research. We would like to thank the many contributors to the 1000 Functional Connectomes Project and INDI; it is their vision and contributions have made these efforts successful. And to the many of members of the INDI team over the years, especially Maarten Mennes, Quiyang Li, Dan Lurie, and David O’Connor. We thank the Neuroimaging Informatics Tools and Resources Clearinghouse (NITRC) for hosting support for INDI, as well as Amazon Web Services and the COllaborative Informatics and Neuroimaging Suite (COINS). This work was supported in part by gifts to the Child Mind Institute from Phyllis Green, Randolph Cowen, Joseph P. Healey and the Stavros Niarchos Foundation. ADM received grant support from the National Institute of Health (521MH107045); XNZ received support from the National Basic Research Program (2015CB351702), National Natural Science Foundation of China (81220108014) and Beijing Municipal Science & Technology Commission (Z161100002616023). Primary funding for the NKI-RS initiatives is provided by grants from the NIH (R01MH094639, R01MH101555, R01- AG047596, U01MH099059), as well as support from the New York State Office of Mental Health and Research Foundation for Mental Hygiene, and Child Mind Institute (1FDN2012-1).

## Author Contributions

Conception and Experimental Design:
AK, ADM, BB FXC, MPM, RCC, SC, XNZ

Data Organization and Scoring:
 AK, AKr, BK, HX, JC, JS, MPM

Data Analysis:
AK, JS, JC, MF, RCC, MPM

Interpretation of Findings:
All authors contributed to the interpretation of findings.

Drafting of the Manuscript:
AK, MPM

Critical Review and Editing of the Manuscript:
All authors contributed to the critical review and editing of the manuscript.

